# Bisphenol A derivatives act as novel coactivator binding inhibitors for estrogen receptor β

**DOI:** 10.1101/2021.05.10.443431

**Authors:** Masaki Iwamoto, Takahiro Masuya, Mari Hosose, Koki Tagawa, Tomoka Ishibashi, Eiji Yoshihara, Michael Downes, Ronald M. Evans, Ayami Matsushima

## Abstract

Bisphenol A and its derivatives are recognized endocrine disruptors based on their complex effects on estrogen receptor (ER) signaling. While the effects of bisphenol derivatives on ERα have been thoroughly evaluated, how these chemicals affect ERβ signaling is not well understood. Herein, we identified novel ERβ ligands by screening a chemical library of bisphenol derivatives. Many of the compounds identified showed intriguing dual activities as ERα agonists and ERβ antagonists. Docking simulations suggested that these compounds act as coactivator binding inhibitors (CBIs). Direct binding experiments using wild-type and mutated ERβ demonstrated the presence of a second ligand interaction position at the coactivator binding site in ERβ. Our study is the first to propose that bisphenol derivatives act as CBIs, presenting a critical view point for future ER signaling-based drug development.

## Introduction

Estrogen receptors (ERs) are members of the nuclear receptor family of transcription factors that directly bind to consensus nucleotide sequences to induce gene transcription. 48 human nuclear receptors have been identified, including those for sex steroid hormones, glucocorticoids, retinoids, and vitamin D (*1, 2*), with many of these receptors recognized as therapeutic targets for a wide range of diseases (*3*). In particular, ERs are major drug targets for breast cancer (*4*) and menopausal disorders. Two ER isoforms exist, ERα and ERβ, that have high amino acid similarity in both the DNA-binding domains (DBDs) and ligand-binding domains (LBDs) (*5*). Many ERα and/or ERβ -associated gene disruption experiments have been reported (*6*). Female mice lacking ERα are infertile, while males exhibit decreased fertility (*7*). Disruption of ERα in female mice leads to hypoplastic uteri, and ERα-disrupted females do not respond to estradiol treatments. ERβ knockout mice present with less severe phenotypes than those with ERα knockout, even though ERβ - disrupted female mice are subfertile predominantly due to reduced ovarian efficiency (*8*). Moreover, ERα and ERβ double-knockout mice show normal reproductive tract development during the prepubertal period. However, those animals present with similar features to ERα knockout mice during adulthood. Furthermore, this diagnostic phenotype indicates that ERβ plays a role in oocyte progression in the postnatal ovary (*9, 10*). Both ERα and ERβ are activated by endogenous estrogens, however, their expression patterns and actions are different (*11*), with each receptor assumed to have specific biological functions.

A growing body of work in laboratory animals supports bisphenol A (BPA) as an endocrine disrupting chemical (EDC) (*12*) that has adverse effects on not only the female reproductive system, but also on the brain and immune system (*13*). BPA is used extensively as a raw material for making polycarbonate plastics and epoxy resins. However, its likely adverse effects on humans, especially infants and fetuses, has recently led to BPA being phased out of polycarbonate plastic and resin production (*14*). Various BPA derivatives have been developed to create more firm and stable plastics and resins, and these derivatives are now preferred as raw materials (*15*) (Fig. 1). However, BPA analogues have already been detected in the environment (*15, 16*). Fluorine-containing BPA, i.e., bisphenol AF (BPAF, 2,2-Bis(4-hydroxyphenyl)hexafluoropropane, CAS No. 1478-61-1), is seen as a practical alternative to BPA, despite reported estrogenic activity in MCF-7 breast cancer cells (*17*). Eight BPA derivatives, including BPAF, have been detected in sediments collected from industrialized areas (*18*) and indoor dust (*19*). In addition, BPA analogs have been found in urine samples from individuals living close to a BPAF manufacturing plant (*20*) and a municipal solid waste incineration plant(*21*). Chlorine-containing BPA, i.e., bisphenol C, (BPC, also known as bisphenol C2 or bisphenol Cl2, 1,1-Dichloro-2,2-bis(4-hydroxyphenyl)ethylene, CAS No. 14868-03-2), is a beneficial substrate for polymer production due to the high thermal stability of BPC-containing polycarbonate (*22, 23, 24*). Notably, BPC is structurally similar to two banned pesticides dichlorodiphenyltrichloroethane (DDT, 1,1′-(2,2,2-trichloroethylidene)bis(4-chlorobenzene), CAS No. 50-29-3) and methoxychlor (1,1′-(2,2,2-trichloroethylidene)bis(4-methoxybenzene), CAS No. 72-43-5) (*25, 26*). Based on its high affinity for endogenous ERs in MCF-7 cells (*27*), BPC was considered but ultimately not included in the list of *in vitro* endocrine disruptors by the Interagency Coordinating Committee on the Validation of Alternative Methods (ICCVAM) (NIH Publication No: 03-4503) in 2003. Historically, the designation of 2,2-bis(4-hydroxy-3-methylphenyl) propane (CAS No. 79-97-0, which does not have chlorine atoms), as BPC has led to some confusion in the literature, however chlorine-containing BPC has been detected in human breast milk (*28*).

**Fig. 1.**
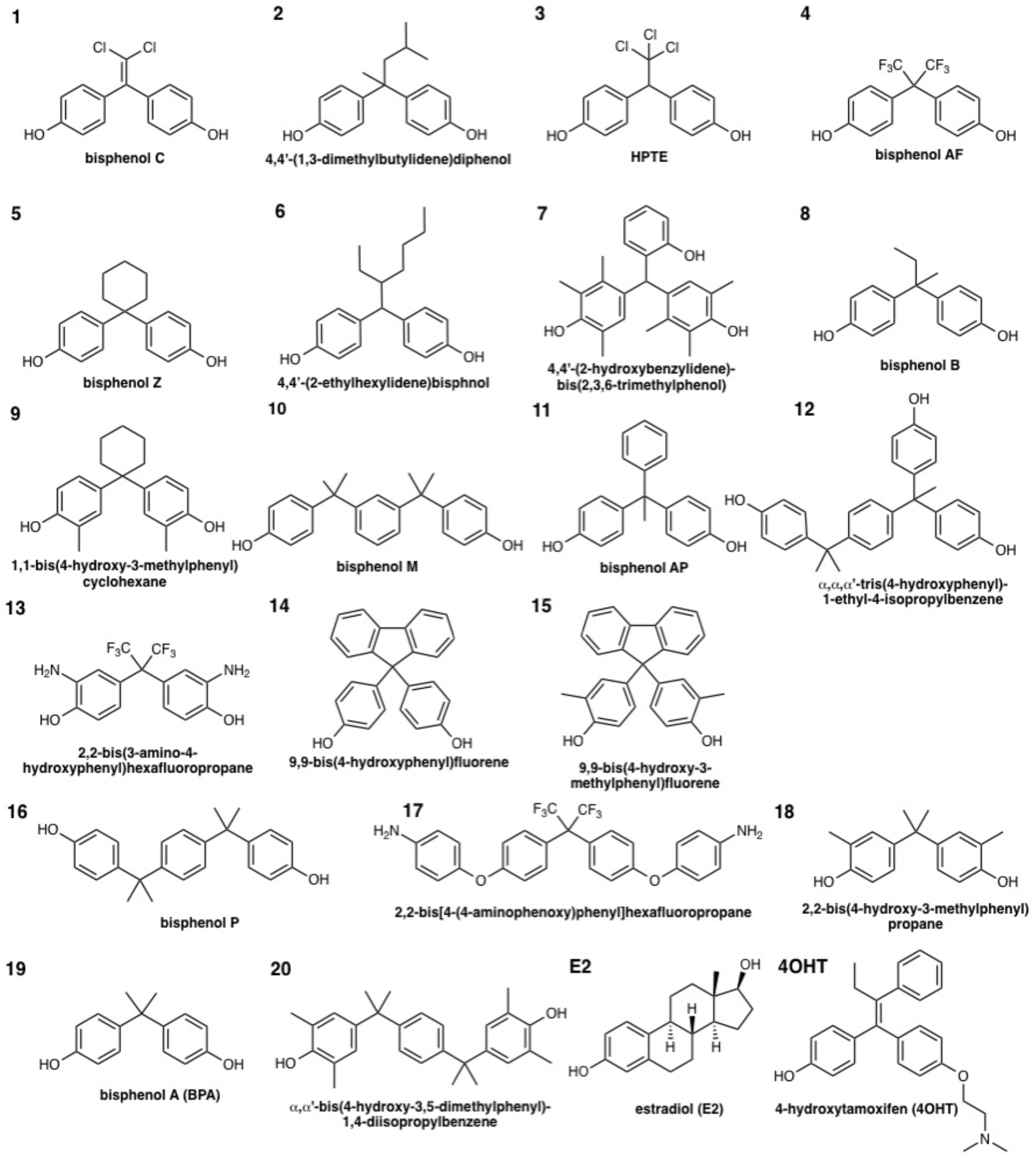
Structures of BPA derivatives selected via screening using an ERβ binding assay. Chemical structures of E2, 4OHT, and 20 BPA-related compounds exhibited stronger binding abilities than, or comparable to, BPA; BPC had the highest binding ability to ERβ. Fluorine-containing BPA derivatives, i.e., 9,9-Bis(4-hydroxyphenyl)fluorine and 9,9-bis(4-hydroxy-3-methylphenyl)fluorene, exerted stronger binding abilities than did BPA.

ERα and/or ERβ are major targets of EDCs which interfere with their estrogen-responsive signaling pathways (*29*). Human ERα and ERβ have almost identical DBDs, differing by only two amino acids, and both receptors bind the same estrogen-response elements in transcriptional control regions. Although ERα and ERβ also have similar LBDs, they have some distinctive features in terms of ligand selectivity and target gene regulation (*30*). Endogenous estrogen, 17-β estradiol (E2), binds to ERα slightly stronger than to ERβ. Similarly, BPA binds ERα with higher affinity than ERβ, although its binding abilities are much weaker than those of E2. In contrast, BPAF and BPC display higher affinity for both ERα and ERβ than BPA, with a preference for ERβ over ERα binding. BPAF and BPC show antagonistic activity against ERβ in reporter gene assays using HeLa cells (*37, 32*). BPAF and BPC show much stronger antagonist activity for ERβ than ERα, (*32, 33*). While crystal structures have provided insight into ERα activation/inactivation mediated by of BPAF and BPC binding (*32, 33*), the structural changes induced by the strong antagonistic activity of BPAF and BPC against ERβ are not well established. Recently, we found that the bisphenol moiety is a privileged structure for ERα. Here we describe the biphasic binding of BPAF and BPC to ERβ and propose a novel two-site binding model of the ERβ - BPC complex, based on the crystal structure of 4-hydroxytamoxfen (4OHT) bound to ERβ. This is the first study to mechanistically associate the antagonistic actions of EDCs with interactions at the coactivator-binding site, thereby providing insight into developing safer raw materials that do not exhibit endocrine-disrupting features.

## Results

### The bisphenol scaffold binds both ERα and ERβ

We screened a library of 119 bisphenol derivatives and related compounds using a radioligand competitive binding assay with [^3^H]E2 for ERβ. Some of these bisphenol derivatives have been detected in human biological samples (*16*). The CAS registry numbers (RNs), common names, and IUPAC names are provided in Supplementary Table 1. We found 18 bisphenol derivatives with similar or stronger ERβ binding compared to BPA (Table 1 and Fig. S1). BPC showed the strongest ERβ (IC_50_: 2.99 nM), and highest ERα (IC_50_ of 2.81 nM) binding affinity of the derivatives examined. The second strongest ERβ binding was seen with Compound No.2 (4,4′-(1,3-dimethylbutylidene)bisphenol; IC_50_: 16.1 nM), although higher affinity was measured with ERα (IC_50_: 5.75 nM). 4,4′-(1,3-Dimethylbutylidene)bisphenol 2,2-bis(*p*-hydroxyphenyl)-1,1,1-trichloroethane (HPTE) (3) and BPAF showed comparable binding ability to ERβ (IC_50_: ~18 nM). Contrary to the results for 4,4′-(1,3-dimethylbutylidene)bisphenol (2), HPTE (3) and BPAF were preferential ERβ ligands, displaying three times stronger binding to ERβ than ERα. Although bisphenol Z (BPZ, 5), 4,4′-(2-ethylhexylidene)bisphenol (6) and 4,4′-(2-hydroxybenzylidene)-bis(2,3,6-trimethylphenol) (7) showed similar results to BPAF, they bound more strongly to ERα. The majority of the chemicals tested elicited comparable binding to both ERα and ERβ. Of the 18 derivatives with similar or stronger ERβ binding compared to BPA, 14 showed slightly stronger binding abilities to ERα than ERβ (Table 1). We reported that 18 bisphenol derivatives bound to ERα more strongly than did BPA (*34*). Bulky functional groups at their sp^3^-carbon connecting two phenol groups were beneficial for ERβ binding, similar to the results previously observed for ERα (*34*). However, ERβ binding abilities did not precisely correlate with those of ERα. Fluorene derivatives, 9,9-bis(4-hydroxyphenyl)fluorene (14) and 9,9-bis(4-hydroxy-3-methylphenyl)fluorene (15), not only bound to ERα (*34, 35*) but also to ERβ (*35*), with their ERβ binding ability stronger than that of BPA. Bisphenol derivatives possessing halogen atoms between two phenol groups, especially chlorine-containing derivatives, showed strong ERβ binding.

**Table 1.** Receptor binding affinity (mean ± SD) of BPA derivatives for ERβ. Receptor binding affinity were evaluated by competitive binding assay using [^3^H] 17β - estradiol as a radioligand.

| Compound No. | Chemicals | Binding affinity (IC<sub>50</sub>, nM) |  |  |  |
| --- | --- | --- | --- | --- | --- |
| | | ERβ | | ERα<sup>[34]</sup> | |
| E2 | estradiol | 2.17 | ± 0.6 | 0.88 | ± 0.13 |
| 1 | bisphenol C | 2.99 | ± 1.0 | 2.81 | ± 0.61 |
| 4OHT | 4-hydroxytamoxifen | 4.66 | ± 1.5 | 2.85 | ± 0.20 |
| 2 | 4,4'-(1,3-dimethylbutylidene)bisphenol | 16.1 | ± 6.1 | 5.75 | ± 1.92 |
| 3 | 2,2-bis(*p*-hydroxyphenyl)-1,1,1- trichloroethane (HPTE) | 18.1 | ± 4.9 | 59.1 | ± 1.5 |
| 4 | bisphenol AF | 18.9 | ± 8.4 | 53.4 | ± 7.3 |
| 5 | bisphenol Z | 21.6 | ± 1.9 | 56.9 | ± 0.6 |
| 6 | 4,4'-(2-ethylhexylidene)bisphenol | 25.9 | ± 8.5 | 18.5 | ± 6.7 |
| 7 | 4,4'-(2-hydroxybenzylidene)-bis(2,3,6-trimethylphenol) | 41.5 | ± 2.0 | 12.3 | ± 7.3 |
| 8 | bisphenol B | 79.8 | ± 12.6 | 195 | ± 44 |
| 9 | 1,1-bis(4-hydroxy-3-methylphenyl)cyclohexane | 132 | ± 6.5 | 38.6 | ± 7.2 |
| 10 | bisphenol M | 148 | ± 80 | 56.8 | ± 11.7 |
| 11 | bisphenol AP | 158 | ± 33 | 259 | ± 41 |
| 12 | $\alpha$, $\alpha$, $\alpha'$-tris(4-hydroxyphenyl)-1-ethyl-4-isopropylbenzene | 212 | ± 36 | 61.7 | ± 10.4 |
| 13 | 2,2-bis(3-amino-4-hydroxyphenyl)hexafluoropropane | 224 | ± 113 | 334 | ± 112 |
| 14 | 9,9-Bis(4-hydroxyphenyl)fluorene | 325 | ± 60 | 2230 | ± 202 |
| 15 | 9,9-bis(4-hydroxy-3-methylphenyl)fluorene | 405 | ± 108 | 321 | ± 103 |
| 16 | bisphenol P | 607 | ± 28 | 176 | ± 35 |
| 17 | 2,2-bis[4-(4-aminophenoxy)phenyl]hexafluoropropane | 609 | ± 81 | 1030 | ± 375 |
| 18 | 2,2-bis(4-hydroxy-3-methylphenyl)propane | 744 | ± 429 | 368 | ± 22 |
| 19 | bisphenol A | 900 | ± 70 | 1780 | ± 764 |
| 20 | $\alpha$, $\alpha'$-bis(4-hydroxy-3,5-dimethylphenyl)-1,4-diisopropylbenzene | 10000 | > | 733 | ± 628 |

To gain insight into the differences observed in ERβ and ERα binding, we compared the ligand binding cavities in the deposited ERβ and ERα LBD crystal structures. The sizes of the canonical binding pockets were calculated for 45 ERα and 25 ERβ structures in their active conformations using SiteFinder function, and the amino acid residues surrounding the bound ligands identified (Tables S2 and S3). The average ERβ pocket was smaller than for ERα, (430.9 Å^3^ and 369.3 Å^3^ for ERα and ERβ, respectively; Fig. 2A). The typical ligand-binding pockets of each receptor in the active conformation is illustrated (Figs. 2C and 2D). Moreover, the average size of the ligand binding pocket in 17β -estradiol-bound ERα and ERβ structures was 419.4 Å^3^ and 385.0 Å^3^, respectively, and in genistein-bound ERα and ERβ structures was 475.9 Å^3^ and 375.8 Å^3^, respectively. Although these results suggested that ERα is able to accept larger ligands than ERβ, the amino acid residues surrounding the ligands differ slightly. Some of the smaller ligands fit more adequately into the ERβ compared to the ERα ligand binding pocket.

**Fig. 2.**
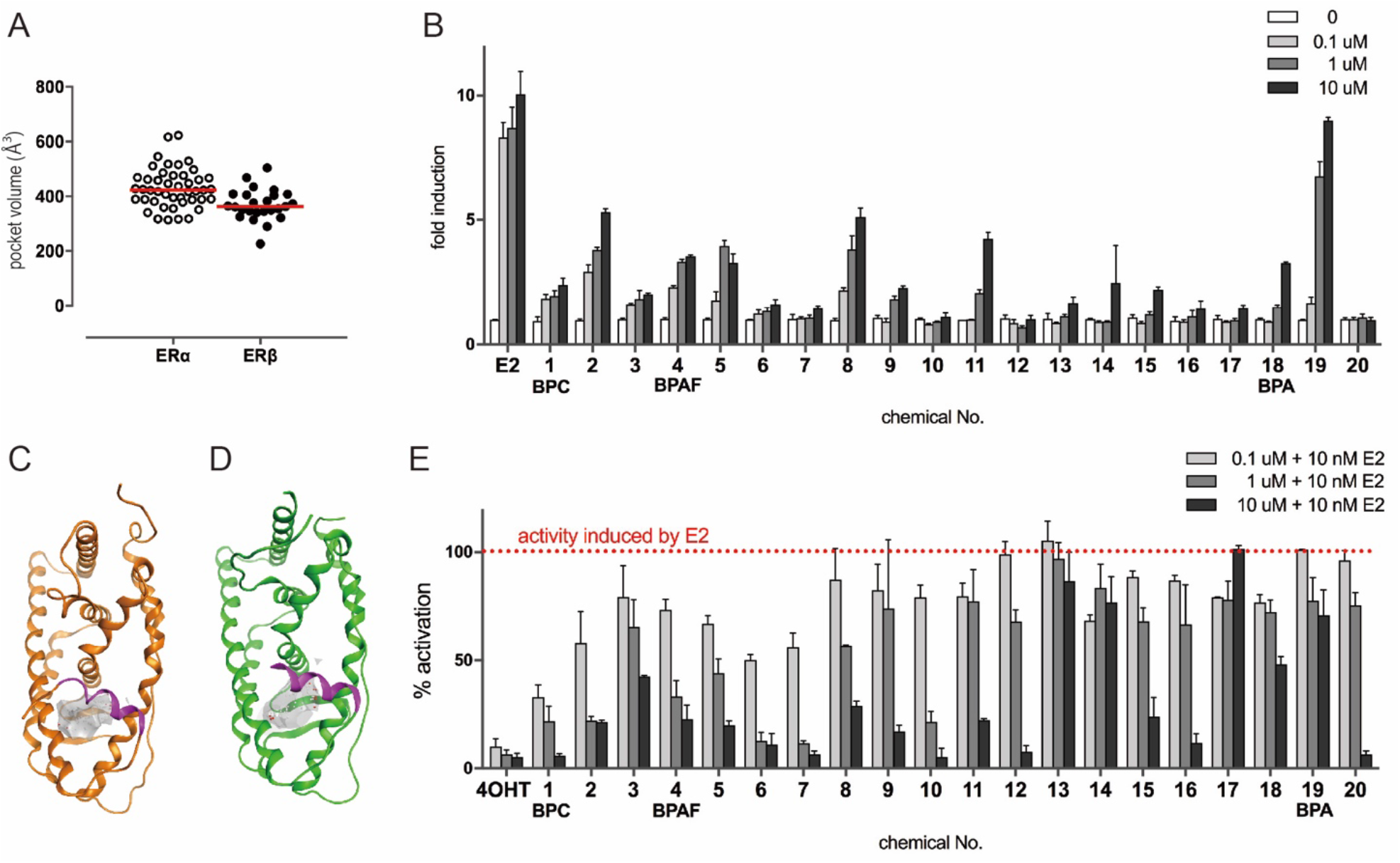
Differential activities of BPA derivatives on ERα and ERβ. (A) Ligand-binding pocket volumes from ERα (open circles) and ERβ (filled circles) calculated from crystal structures in the presence of activating ligands; average volumes indicated by red lines. (B) Top 10 BPA derivatives binding to ERβ induced partial agonistic activity against ERβ. (C) The ligand-binding pockets of ERα (PDB ID 1QKU) and (D) ERβ (3OLL) are illustrated in gray; estradiol is bound as the ligand. (E) Sixteen chemicals, including tricyclic bisphenols, inhibited more than half of the 10 nM E2-induced transcriptional activity.

### BPC and BPAF bind but fail to activate ERβ

Reporter assays using HeLa cells were performed to evaluate ERβ transcriptional activity induced by BPA, BPC, BPAF, and 17 bisphenol derivatives (Fig. 2B). BPA elicited the strongest ERβ agonistic activity of the derivatives, with the activity at 10 μM comparable to that seen with the endogenous ligand E2 despite its affinity being 400 times weaker than that of E2. 4,4′-(1,3-dimethylbutylidene)bisphenol (2) and bisphenol B (8) achieved ~50% of BPA-induced transcriptional activity at the highest concentration of 10 μM. While compound 2, found as an impurity in industrial-grade BPA, has been shown to function as an ERα agonist in yeast-two hybrid assays (*36*), our results reveal a high affinity for and functional activation of ERβ. Compound 2 and 8 are structurally similar to BPA, possessing one methyl group on the sp^3^-carbon that bridges the two phenol groups, suggesting that this conformation is beneficial for ERβ activation. BPC, HTPE, BPAF, BPZ, 1,1-bis(4-hydroxy-3-methylphenyl)cyclohexane (9), 9,9-bis(4-hydroxy-3-methylphenyl)fluorene (15), and 2,2-bis(4-hydroxy-3-methylphenyl)propane (18) functioned as partial agonists, inducing 20% to 30% of the E2-induced transcriptional activity. The transcriptional activity of BPC, HPTE, and BPAF was consistent with a previous report investigating ERα and ERβ, in which these compounds elicited weaker activity against ERβ than ERα (*32, 33*). Surprisingly, 4,4′-(2-ethylhexylidene)bisphenol (6), 4,4′-(2-hydroxybenzylidene)-bis(2,3,6-trimethylphenol) (7), bisphenol M (10), α, α, α′-tris(4-hydroxyphenyl)-1-ethyl-4-isopropylbenzene (12), bisphenol P (16), and α,α′-bis(4-hydroxy-3,5-dimethylphenyl)-1,4-diisopropylbenzene (20) showed no agonist activity against ERβ. These findings contrast with ERα, where the majority of bisphenol derivatives with strong binding affinity also showed strong agonistic activity (*34*).

### BPA derivatives function as ERβ antagonists

The finding that many BPA derivatives with high binding affinities showed almost no agonist activity suggested that they function as ERβ antagonists. To explore this possibility, the inhibitory effects of the BPA derivates (100 pM, 1 μM, 10 μM) against 10 nM E2-induced ERβ activation were measured (Fig. 2E). BPC showed the strongest antagonistic activity, with additional halogen-containing bisphenols (i.e., HPTE, and BPAF) also elicited antagonistic activities, consistent with previous reports (*31-33*). 4,4′-(1,3-dimethylbutylidene)bisphenol (2), which had the second strongest binding ability and partial agonist activity compared to BPA, showed weak antagonist activity, contrasting with its reported ERα agonism. Bisphenol B (8) showed similar weak antagonist activity, with both Bisphenol B (8) and 4,4′-(1,3-dimethylbutylidene)bisphenol (2) inhibiting 50% of BPA-induced activation. Tricycle bisphenols (i.e., bisphenol M (10), α, α, α′-tris(4-hydroxyphenyl)-1-ethyl-4-isopropylbenzene (12), bisphenol P (16), and α,α′-bis(4-hydroxy-3,5-dimethylphenyl)-1,4-diisopropylbenzene (20)) showed antagonistic activity, presumably through the disruption of the active conformation, as reported for ERα (*34*). While demonstrating no agonist activity, 4,4′-(2-ethylhexylidene)bisphenol (6) and 4,4′-(2-hydroxybenzylidene)-bis(2,3,6-trimethylphenol) (7), suppressed 90% of E2-induced activation at the 10 μM concentration. Interestingly, the fluorene derivative, 9,9-bis(4-hydroxy-3-methylphenyl)fluorene (15) functioned as a weak antagonist, demonstrating that fluorene derivatives 14 and 15 can exhibit both ERβ and ERα antagonistic activity (*34, 35*). With the exception of the tricyclic bisphenols, these findings indicate that most bisphenol derivatives with strong ERβ binding functioned as antagonists, even though they showed only agonist activities to ERα (*34*).

### Docking analysis predicts BPC binding to the surface of ERβ

To investigate the contrasting actions of BPA derivatives as ERβ antagonists and ERα agonists, we performed docking simulations using the LBD of human ERβ and BPC, the strongest binder among the BPA derivatives examined using a competitive binding assay with [^3^H]E2. Possible ligand binding sites in 38 deposited ERβ crystal structures were identified using SiteFinder, a program for binding site analysis equipped in the Molecular Operating Environment (MOE). Canonical, as well as putative binding sites were ranked according to PLB, a specific parameter in SiteFinder (*37*). Consistently, the top five predicted sites in each structure were the canonical ligand-binding sites. Interestingly, an actual surface 4OHT binding site close to the hydrophobic groove for the coactivator recognition surface of ERβ (PDB ID: 2FSZ) was ranked 11^th^ in the PLB order. Moreover, this location was a predicted binding site on all antagonist-bound ERβ structures, based on PLB. Notably, this second site was not predicted as a binding site on over half of the agonist-bound structures (Supplementary Table S4). These predictions suggest that ERβ antagonism induced by BPC and other BPA derivatives may be due to inhibition of coactivator recruitment. Next, we performed a docking simulation for ERβ LBD and BPC using both its canonical and second binding sites as target rooms. BPC was able to fit and bind in both rooms, with one of its chlorine atoms interacting with the tryptophan residue (Trp335) on helix 5 via halogen interaction (Fig. 3, A and B). The obtained model structure suggested that BPC binding to the second binding site prevented recruitment of coactivators for gene transcriptions, similar to 4OHT (Fig. 3, C and D). We hypothesized that the binding affinity of BPA derivatives to this coactivator binding site would correlate with antagonistic activity. To explore this notion, docking simulations were performed for each BPA derivatives (Fig. S2) and the free energy of ligand binding evaluated using a docking simulation and the GBVI/WSA dG scoring function (larger negative scores indicate more stable ligand/receptor complexes) (*38*). Correlation of the GBVI/WSA dG scores with the extent of antagonism (reported as the % inhibition of 10 nM E2 induced transcriptional activity) revealed a linear relationship (correlation coefficient of − 0.83), suggesting that inhibition of coactivator recruitment underlies the antagonism of ERβ by BPA derivatives (Fig. 3E).

**Fig. 3.**
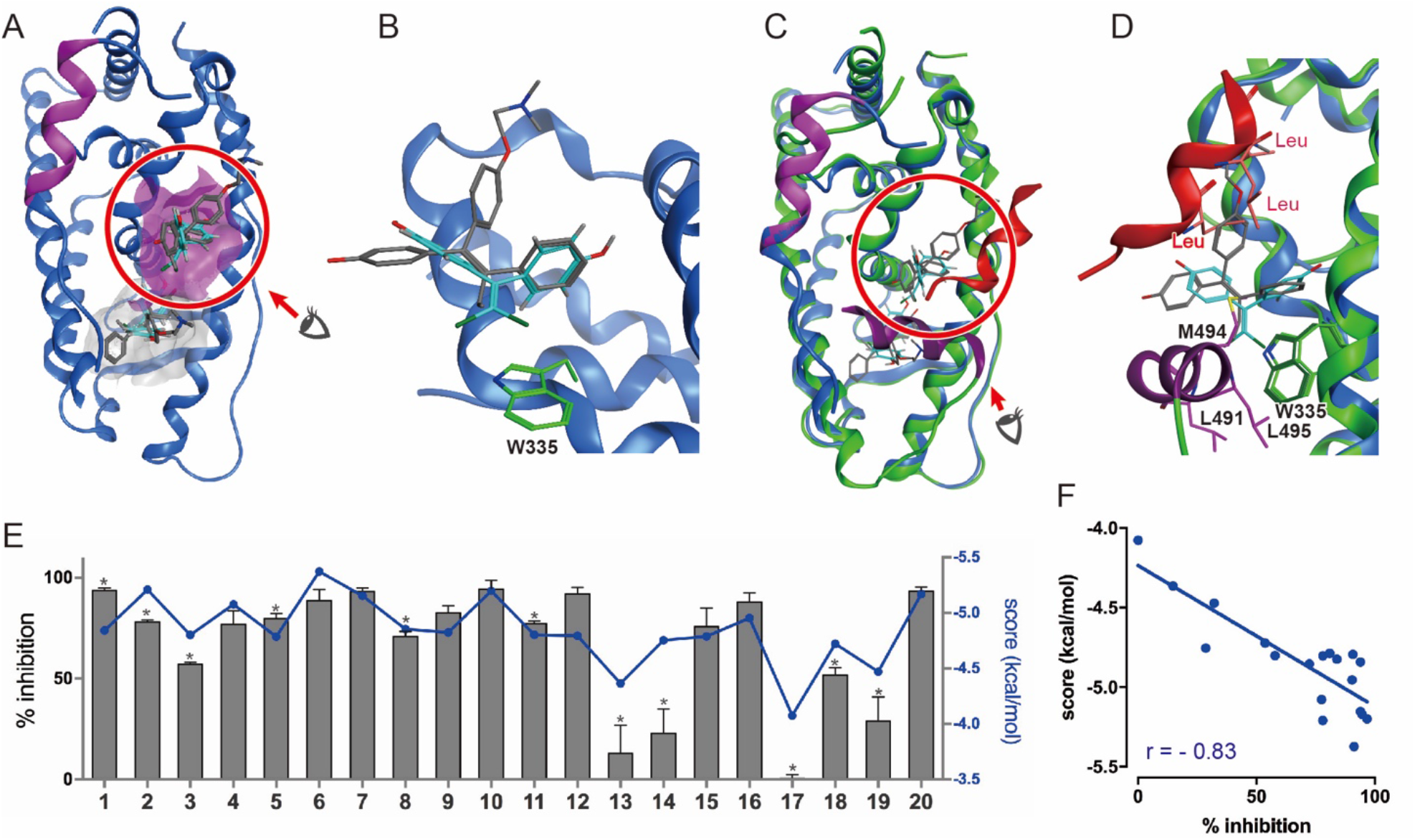
ERβ harbors two ligands in its LBD. (A) Two BPC bound to ERβ during the docking simulation. The canonical binding site is indicated in gray; the second binding site, located on the surface of the receptor, is shown in magenta. The activation helix, H12, is indicated in magenta. (B) Chlorine, a halogen atom of BPC, interacted with the Trp335side chain via halogen interaction in the second binding site. BPC and 4OHT are illustrated in blue and gray, respectively, in the stick model. (C) Superimposition of the calculated BPC-bound ERβ structure (blue) and its agonist form with the nuclear receptor coactivator 1, SRC1. (green, PDB ID, 3OLL). SRC1 is indicated as a red α-helix, H12 of its agonist form is indicated in purple, BPC is illustrated in blue, and 4OHT is shown in gray. BPC clashed with the amino acid residues on H12 in the ERβ agonist form; therefore, BPC prevented the ERβ activation. BPC and 4OHT disrupted the SRC1 binding due to steric hindrance of the amino acid residues shown in the red stick models. (D) In ERβ -agonist form, amino acid residues surrounding Trp335 within 4.5 Å on H12 are shown in the purple stick model, while leucine residues on the SRC1 LXXLL motif are indicated via the red stick model. (E, F) Correlation of the calculated binding scores and inhibitory activity for ERβ. Inhibitory activity is defined as the ratio of chemicals inhibiting transcriptional activity induced by 10 nM E2. \**p*< 0.05.

### Biphasic 4OHT binding indicative of two ERβ binding sites

To further support the presence of a second ligand binding site, competitive binding assays were performed using BPA, BPC and BPAF and tritium-labeled 4OHT ([^3^H]4OHT) (Fig. 4A). Notably, a biphasic dose-response curve was observed for BPC (18.1 nM and 2281 nM IC_50_) that was not evident in the [^3^H]E2 competitive analyses. Similarly, BPAF displayed a biphasic binding curve, albeit with weaker binding at both the high- and low-affinity sites compared to BPC. Moreover, 4OHT showed a biphasic curve, consistent with the 4OHT/ERβ crystal structure (PDB:ID 2FSZ). In contrast, BPA, which did not elicit antagonistic activity, showed a sigmoidal curve indicative of a single ligand binding site. Interestingly, the tri-fluorine substitution of the methyl groups in BPAF increased ERβ binding ~50 fold compared to BPA. These results confirmed the presence of two distinguishable binding sites for BPC and BPAF on ERβ. In contrast, the typical sigmoidal curves seen in E2 competitive binding assays using [^3^H]4OHT and [^3^H]E2 are indicative of single ligand binding site.

**Fig. 4.**
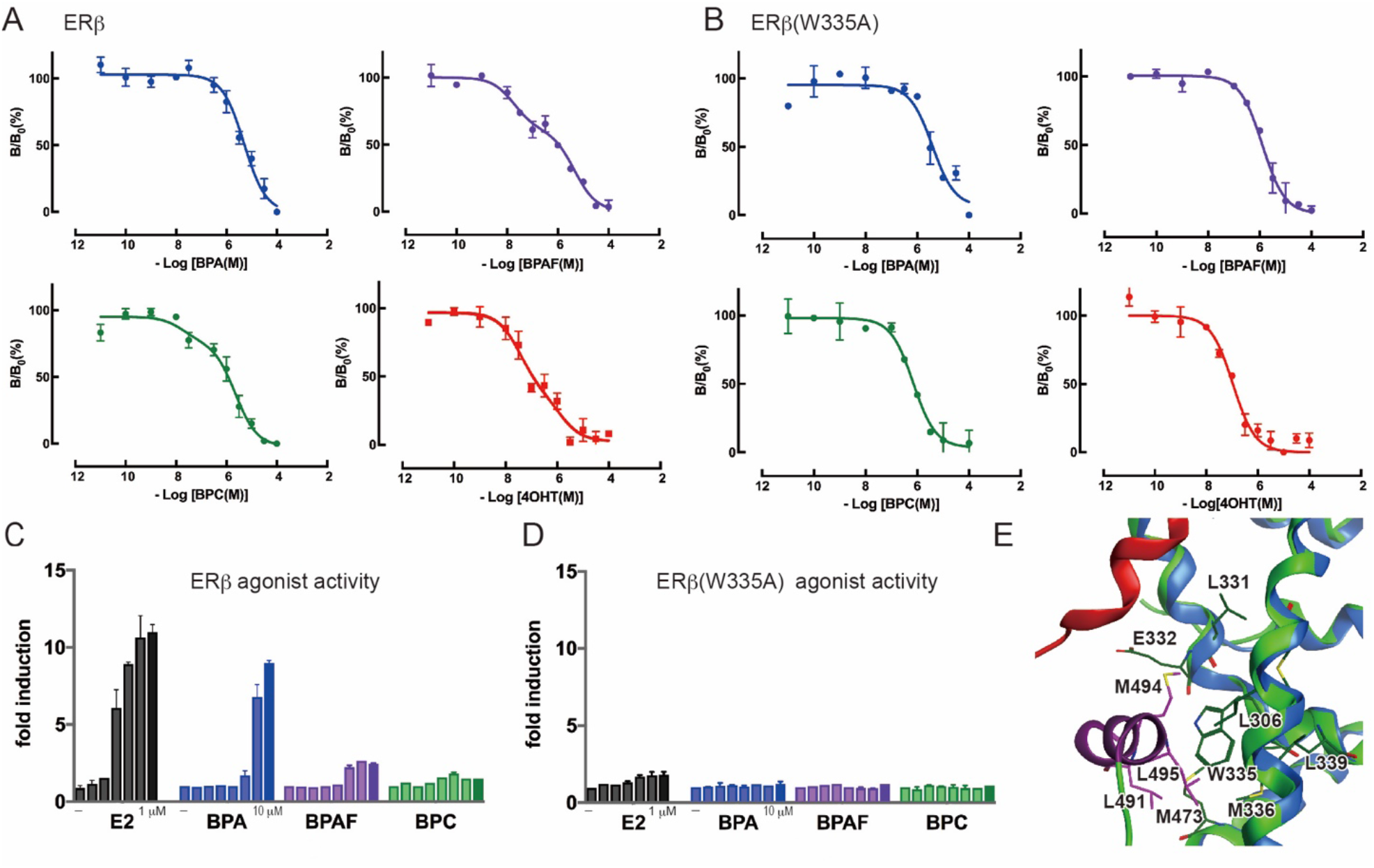
Binding properties and transcriptional features of BPAF and BPC showed the importance of ERβ W335 for their receptor binding and activation. (A) Detailed competitive binding curves of BPA, BPAF, BPC, and 4OHT using [^3^H]4OHT illustrated a diphasic binding curve, in which chemicals compete with [^3^H]4OHT in two binding sites on wild type ERβ. (B) ERβ (W335A) competitive binding assays showed typical sigmoidal binding curves. (C) The reporter gene assay indicated that BPAF and BPC induced weak transcriptional activity in wild type ERβ, while E2 and BPA showed strong transcriptional activity. (D) ERβ (W335A) lost E2 or BPA-induced transcriptional activity, indicating that Trp335 substitution disrupted active conformation. (E) In ERβ agonist form, amino acid residues surrounding Trp335 within 4.5 Å are represented as green and purple stick models. (PDB ID, 3OLL).

### Trp335 is required for biphasic ligand binding

The docking simulations suggested that hydrophobic interactions between the BPA derivatives and the indole group of Trp335 were required for ERβ binding, and identified a potential halogen interaction between the chlorine atom of BPC and the indole ring. To determine the contributions of these putative interaction to BPC binding, the corresponding tryptophan was mutated to alanine (A). Saturation binding assays revealed a typical sigmoidal dose-response curve and a *K*_d_ of 23.1 nM for E2 against ERβ (W335A), indicating preservation of the canonical binding site (Fig. S3, A).

Competitive binding assays confirmed two 4OHT binding sites in ERβ, with *K*_d_ values of 4.6 nM and 53.1 nM. In contrast, a single binding site was evident in ERβ (W335A) (*K*_d_ 34.2 nM) (Fig. S3, B). Similarly, the biphasic binding of BPC and BPAF were lost in the ERβ (W335A) mutant (Fig. 4, A and B). The IC_50_ values of 4OHT, BPC, and BPAF were 106 ± 51 nM, 691± 29 nM, and 1249 ± 579 nM, respectively. BPA illustrated a typical sigmoidal competitive dose-response curve against ERβ (W335A), similar to the result against ERβ. These results indicated that replacing Trp for Ala compromises the second 4OHT and BPA derivatives binding site on the surface of ERβ LBD.

### W335A reduces ERβ transcription activity

Reporter assays revealed that E2-induced transcriptional activation was markedly reduced by the tryptophan to alanine substitution in ERβ (Fig. 4, C and D). Given that E2 binding ability was retained, this is consistent with reduced coactivator binding. Indeed, in the active conformation, Trp335 interacts with Leu491, Met494, and Leu495 on H12 (Fig. 4E). These results indicated that Trp335 on the ERβ coactivator-binding site plays an important role, not only in interacting with bisphenol derivatives, but also in recruiting coactivators on the surface of ERβ by stabilizing H12 in its active conformation.

## Discussion

Here we report the ERβ transcriptional activities of BPA derivatives including BPC and BPAF using a combination of receptor binding and reporter assays. Unexpectedly, our results clearly showed that many BPA derivatives function as ERβ antagonists, contrasting with their previously reported ERα agonism. Docking simulations indicated that BPA derivatives bind to a second site located near the coactivator binding site on the surface of ERβ -LBD that requires interactions with Trp335. Mutation of tryptophan to alanine led to the loss of this low affinity binding site in ERβ. These results indicated that some BPA derivatives act as antagonists, although most of endocrine-disrupting chemicals, including BPA, are assumed ER agonists. We previously reported that most of the BPA derivatives examined in this study act as weak agonists for ERα. The results obtained in this study demonstrate the importance of screening for both agonist and antagonist activity, especially against ERβ.

We previously reported that tricyclic bisphenols, i.e., Bisphenol M, α, α, α′-tris(4-hydroxyphenyl)-1-ethyl-4-isopropylbenzene, bisphenol P, and α,α′-Bis(4-hydroxy-3,5-dimethylphenyl)-1,4-diisopropylbenzene, act as antagonists against ERα because of the steric hindrance caused by the third aromatic ring structure (*34*). This study showed that this feature is also valid for ERβ; tricyclic bisphenols act as antagonists not only for ERα but also ERβ. In addition to tricyclic bisphenols, many BPA derivatives, including BPAF and BPC, elicit antagonist activity. Our finding for BPAF and BPC are consistent with reports that both chemicals showed partial agonism for ERα and antagonism for ERβ (*31, 32, 39, 40*).

Several ERα- or ERβ -specific agonists have been reported, including propylpyrazole triol (PPT) that selectively binds to and transcriptionally activates ERα (*41*). The first chemical shown to function as an ERα agonist and ERβ antagonist is 2,2-bis(*p*-hydroxyphenyl)-1,1,1-trichloroethane (HPTE), a metabolite of the banned pesticide, methoxychlor [1,1,1-trichloro-2,2-bis(4-methoxyphenyl)ethane] (*42, 43*). Accumulated knowledge gained from protein crystal structures emphasize the importance of halogens in receptor-ligand interactions (*44, 45*). We found that in addition to the halogen containing BPAF and BPC, many BPA derivatives display ERα agonist activities similar to HPTE. These results indicate the complexity of establishing the mechanisms of action of environmental chemicals that activate or suppress the physiological functions of one or more nuclear receptors. In particular, antagonist activities might be overlocked if both binding affinity and transcriptional activity are not determined, as environmental chemicals are typically categorized based on the ability to active ERs.

Recent studies have indicated the value of small molecules that bind to coactivator protein-binding sites on nuclear receptors (*46*). Coactivator-binding inhibitors (CBIs) have been developed for ERs, an androgen receptor, a progesterone receptor, a vitamin D receptor, a thyroid hormone receptor, a pregnane X receptor, a retinoid X receptor, and peroxisome proliferator-activated receptors (*47–50*). This study is the first to conclude that endocrine-disrupting chemicals can function as CBIs for ERβ, indicating the importance of assessing both agonist and antagonist activities of these chemicals.

In summary, we showed that tricyclic bisphenols elicit antagonistic activity against both ERα and ERβ. Our results also indicate that many next-generation bisphenols are agonists and antagonists of ERα and ERβ. Mutagenesis of an ERβ surface amino acid indicated that these next-generation bisphenols act as CBIs. While *in silico* docking analyses support this mechanism of action, future crystallographic studies will be required to provide more direct information on CBIs. This study highlights the mechanistic complexity of the next-generation of bisphenols acting as endocrine-disrupting chemicals.

## Materials and Methods

### Chemicals

17β-estradiol (E2, CAS RN 50-28-2, >98.9%) was obtained from of Research Biochemicals International (Natick, MA, USA). 4-hydroxytamoxifen (4OHT, CAS RN 68047-06-3, >98%) and 2,2-bis(*p*-hydroxyphenyl)-1,1,1-trichloroethane (CAS RN 2971-36-0, >98.9%) were obtained from Sigma-Aldrich Inc. (St. Louis, MO, USA). 4,4′-dihydroxydiphenylmethane (bisphenol F or BPF, CAS RN 620-92-8, >99.0%) and hexestrol (CAS RN 84-16-2, >99.0%) were obtained from FUJIFILM Wako Pure Chemical Corporation (Osaka, Japan); the remaining 117 chemicals were purchased from Tokyo Chemical Industry Co., Ltd. (Tokyo, Japan). Dimethyl sulfoxide (DMSO), used to dissolve each compound in a 10 mM stock solution, was obtained from Sigma-Aldrich. Tritium-labeled 17*β*-estradiol ([^3^H]E2, 4458 GBq/mmol) and 4-hydroxytamoxifen ([^3^H]4OHT, 2960 GBq/mmol) were purchased form PerkinElmer (Waltham, MA, USA).

### ERβ expression and purification

The LBD of ERβ (amino acids 263-530) was expressed as a glutathione *S*-transferase (GST)-fused protein for receptor binding assays. Human ERβ cDNA was obtained from OriGene Technologies (Rockville, MD, USA). The cDNA of ERβ-LBD was amplified using PCR, and subcloned into an pGEX-6p-1 expression vector (Cytiva, Marlborough, MA, USA). The expression of GST-fused ERβ-LBD was induced by 1 mM isopropyl-β-D-1-thiogalactopyranoside (IPTG) in *Escherichia coli* BL21α at 16°C for overnight. The resulting crude protein was affinity-purified using Glutathione-Sepharose 4B (Cytiva), followed by gel filtration in a Sephadex G-10 column (Cytiva).

### Radioligand binding assay

Radioligand binding assays for ERβ and ERβ(W335A) were performed mainly according to a previously reported method (*31, 34*). Saturation binding assays were conducted with [^3^H]E2 or [^3^H]4OHT using GST-ERβ-LBD or GST-ERβ(W335A)-LBD to evaluate the binding ability of radio-labeled compounds. The reaction mixtures of each LBD (20 ng) and a series of concentrations of [^3^H]E2 (0.01–10 nM) or [^3^H]4OHT (0.1–30 nM) were incubated in a total volume of 100 μL of binding buffer (10 mM Tris-buffered saline (pH 7.4), 1 mM ethylene glycol-bis (2-aminoethylether)-*N, N, N*′, *N*′-tetraacetic acid (EGTA), 1 mM ethylenediaminetetraacetic acid (EDTA), 10% glycerol, 0.5 mM phenylmethylsulfonyl fluoride, 0.2 mM leupeptin, and 1 mM sodium vanadate (V)) at 20°C for 2 h, to analyze total binding. Corresponding reaction mixtures, containing 10 μM non-labeled E2 or 4OHT, were incubated to detect each non-specific binding. [^3^H]E2 or [^3^H]4OHT-specific binding was evaluated by subtracting the obtained radioactivity values of total binding from the those of non-specific binding. Following successive incubation with 100 μL of 0.4% dextran-coated charcoal (DCC) (Sigma-Aldrich) in phosphate-buffered saline (pH 7.4) on ice for 10 min, free radioligands bound to DCC were removed using a vacuum filtration system with a 96-well filtration plate (MultiScreenHTS HV, 0.45-mm pore size, Merck KGaA, Darmstadt, Germany) for the bound/free (B/F) separation. The radioactivity of each eluent was measured using a liquid scintillation counter (LS6500; Beckman Coulter, Fullerton, CA, USA) and Clear-sol I (Nacalai Tesque Inc., Kyoto, Japan).

Calculated specific binding of [^3^H]E2 was assessed using Scatchard plot analysis (*51*). Competitive binding assays were performed to evaluate the binding abilitiy of each test compounds using [^3^H]E2, for a library screening or detailed BPA binding assay. Each compound was dissolved in DMSO to prepare a 1.0 mM stock solution, and further diluted to prepare a serial dilutions (10^−12^M to 10^−5^ M) in binding buffer. To assess their binding abilities, each compound was incubated with GST-ERβ-LBD or GST-ERβ(W335A)-LBD (20 ng) and radio-labeled ligand (5 nM of [^3^H]E2 or 5 nM of [^3^H]4OHT, final concentration) for 2 h at 20°C. B/F separation was performed as described above, and the radioactivity was determined using a MicroBeta microplate counter (PerkinElmer Inc.). The IC_50_ value of each test compounds was calculated from the dose-response curves generated via nonlinear regression analysis using Prism software (GraphPad Software Inc., La Jolla, CA, USA).

### Luciferase reporter gene assay

Transcriptional activities of ERβ and ERβ (W335A) were measured as previously reported previously (*31, 34*). HeLa cells were maintained in Eagle’s minimum essential medium (EMEM) (Nissui Pharmaceutical Co. Ltd, Tokyo, Japan) supplemented with dextran-coated charcoal treated fetal bovine serum (DCC-FBS, 10%, v/v) with at 37°C under 5% CO2. To evaluate agonistic activity, HeLa cells were seeded at a density of 5 × 10^5^ cells per 60-mm dish and cultured for 24 h, followed by transfection of the reporter plasmid (3 μg, pGL4.23/3×ERE) and each expression plasmid (1 μg, pcDNA3.1/ERβ or pcDNA3.1/ERβ (W335A)) using Lipofectamine LTX with Plus Reagent (Thermo Fisher Scientific, Inc.), according to the manufacturer’s instructions. After incubation for 24 h, cells were harvested and seeded onto 96-well plates at 5 × 10^4^ cells/well, and then treated with a series of the test compounds (10^−12^ M to 10^−5^ M, final concentration) diluted with 1% bovine serum albumin/PBS (v/v). After a 24-h incubation, luciferase activity was measured using the ONE-Glo™ Luciferase Assay System (Promega Co., Madison, WI, USA) on an EnSpire multimode plate reader (Perkin Elmer, Inc.). To analyze antagonistic activity, serial concentrations of test compounds (10^−12^ M to 10^−5^ M) were treated in the presence of 10 nM E2, which normally induces full transcriptional activity levels in transiently expressed ERβ.

### Docking simulation of each antagonist onto the ERβ LBD

Three-dimensional (3D) coordinates of the compounds were obtained from the Cambridge Structural Database (CSD-Core, The Cambridge Crystallographic Data Centre, Cambridge, UK). Ligand IDs of compounds utilized for docking simulations are summarized in Supplementary Table S5. For the compounds with no corresponding entry in the CSD-System, 3D coordinates were constructed *in silico* using Gaussian 16 (Gaussian, Inc., Wallingford CT, USA), with the basis set of 6–31G. Docking simulations for the ligand/ERβ complex were performed using a Dock functions in the MOE package (Chemical Computing Group, Montreal, QC, Canada); the free energy of each complex was evaluated according to its GBVI/WSA dG score (*38*). Ligand-binding cavity volumes of the deposited crystal structures were analyzed and calculated using the SiteFinder function in MOE.

## Statistical analysis

Significance of the data between experimental groups was determined using unpaired *t*-tests. Data are presented as the mean ± standard deviation (SD), and P values are presented in each figure legend.

## Supporting information

This article contains supporting information (Figs. S1 to S3 and Tables S1 to S5).

## Acknowledgments

We appreciate R.T. Yu, and A.R. Atkins (Salk Institute for Biological Studies) for their helpful suggestions and discussions. We appreciate Y. Shimohigashi (Kyushu University) for providing the chemical library, and X. Liu (Kyushu University) for providing the ERβ- mutated plasmid. We thank the RIKEN BRC Cell Bank, and Cell Resource Center for Biomedical Research Institute of Development, Aging and Cancer, Tohoku University, for providing the HeLa cells.

## Author contributions

A.M. conceived and designed the experimental approaches. A.M., M.I., and T.M. performed most of the experiments. M.H. contributed to the docking simulation analysis. K.T. and T.I. performed the experiments. A.M. wrote the manuscript. E.Y., M.D., and R.M.E. provided critical comments and contributed to the editing of the manuscript.

## Funding and additional information

This work was supported by JSPS KAKENHI JP17H01881, JP18K19147, 18KK0320, and 20H00635, to A.M, and in part by a grant from Izumi Science and Technology Foundation. R.M.E. holds the March of Dimes Chair in Molecular and Developmental Biology at the Salk Institute, and is supported by the NOMIS Foundation - Science of Health. M.D. and R.M.E. was supported in part by the National Institute of Environmental Health Sciences of the National Institutes of Health under Award Number P42ES010337. The content is solely the responsibility of the authors and does not necessarily represent the official views of the National Institutes of Health.

## Conflict of interests

Authors declare no competing interests.

## Data and materials availability

All data needed to evaluate the conclusions in the paper are present in the paper and/or the Supplementary Materials.

## Supplementary Materials for

**Fig. S1.**
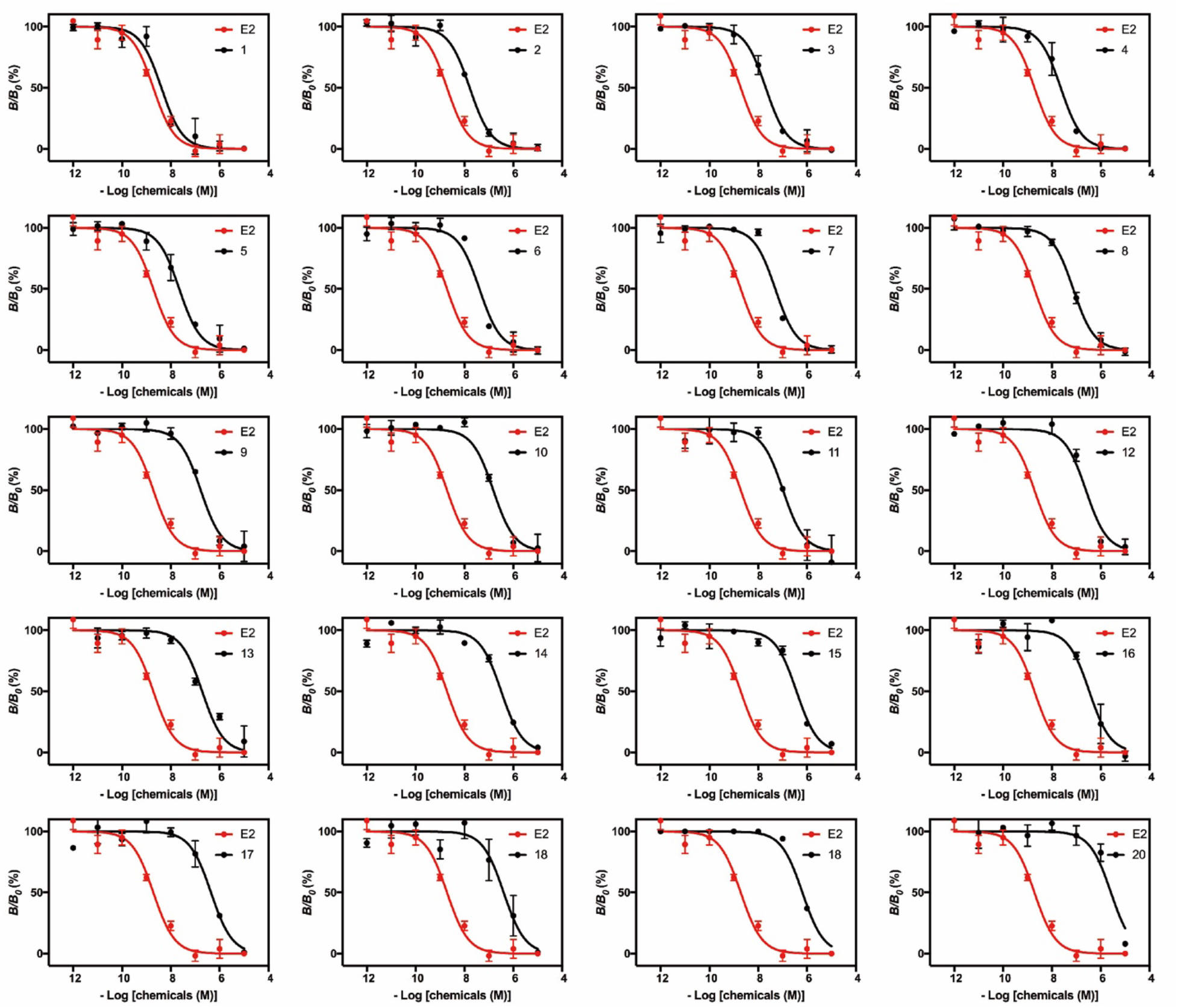
Binding curves indicated the binding ability of each bisphenol derivatives using competitive binding assays with[^3^H]E2. *B*/*B*_0_ is the ratio of displacement by the chemical tested (*B*) against the maximum specific binding (*B*_0_ = 100%) of [^3^H]E2.

**Fig. S2.**
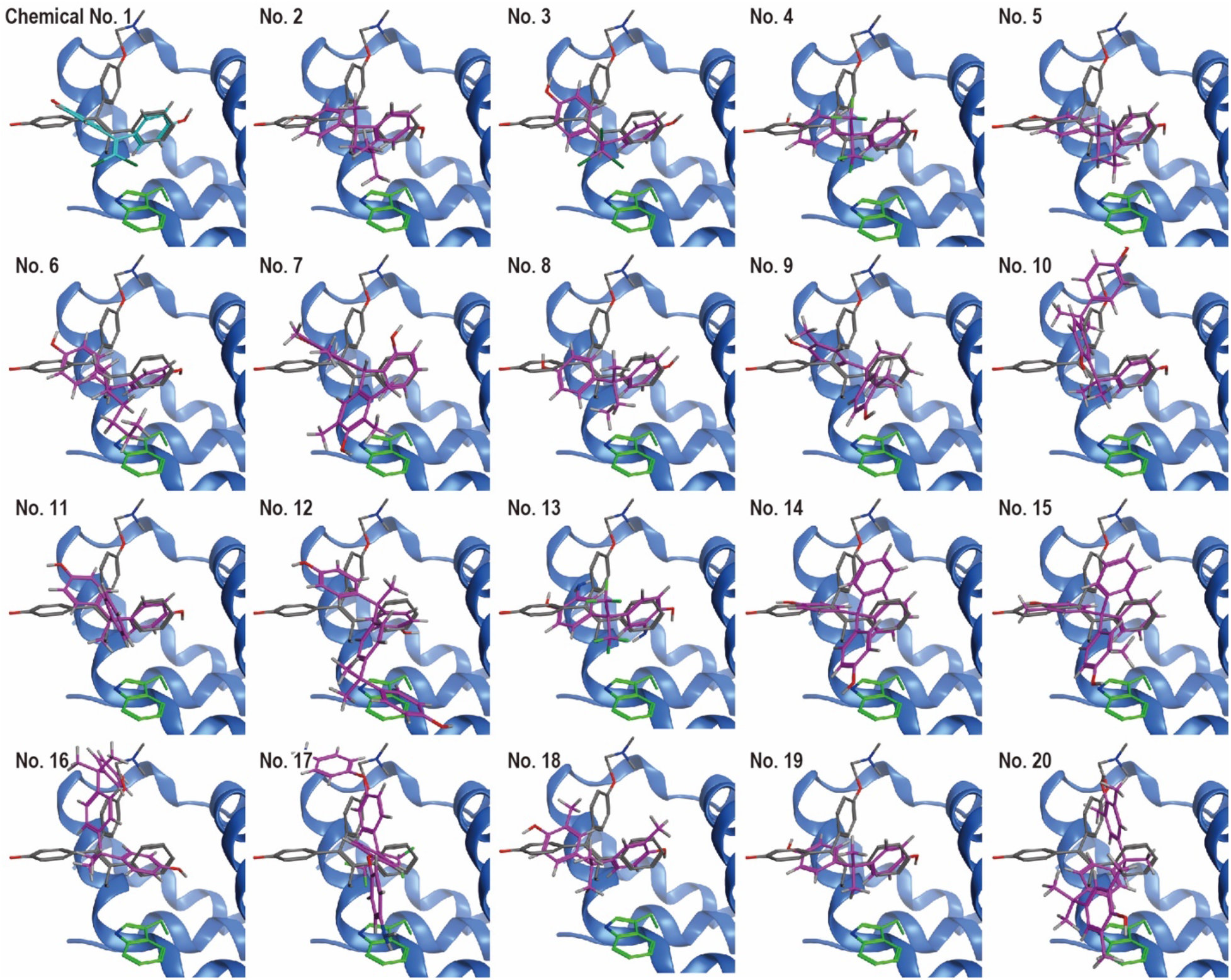
Docking simulation indicated the binding possibilities of BPA derivatives on the second binding site located on the coactivator-binding surface of ERβ. Calculated coordinates of BPC (blue stick model, No. 1) and the other BPA derivatives (magenta stick model, No. 2 to 20) were located close to W335. 4OHT in the crystal structure used for the docking simulation is indicated via a gray stick model.

**Fig. S3.**
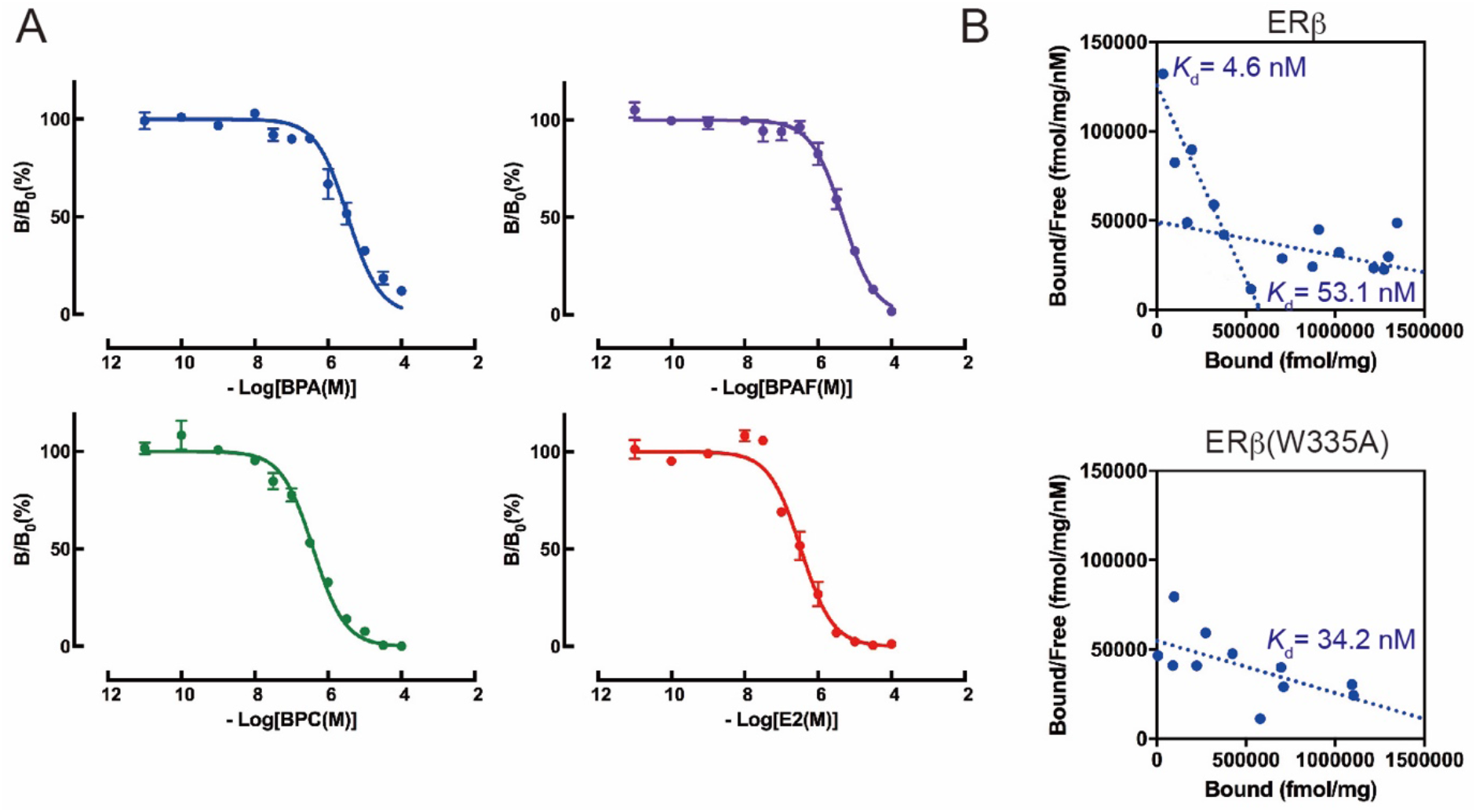
Binding experiments showed that ERβ has two 4OHT binding sites, and only a single binding site for E2. (A) Competitive binding assay of ERβ (W335A) using [^3^H]E2 showed that ERβ (W335A) retained its binding activity for E2 and other BPA derivatives. (B) Saturation binding assays using [^3^H]4OHT estimated that ERβ has both a high and low binding sites for 4OHT, while ERβ (W335A) has only one binding site.

**Table S1.** CAS RNs, common names, and IUPAC names of all the chemicals whose binding ability to ERβ was analyzed using competitive binding assays in this study.

| CAS RN<sup>®</sup> | common name | IUPAC name |
| --- | --- | --- |
| 603-44-1 | 4,4',4''-trihydroxytriphenylmethane | 4,4',4''-methanetriyltriphenol |
| 79-95-8 | tetrachloro bisphenol A | 4,4'-(propane-2,2-diyl)bis(2,6-dichlorophenol) |
| 79-94-7 | tetrabromo bisphenol A | 4,4'-(propane-2,2-diyl)bis(2,6-dibromophenol) |
| 77-40-7 | bisphenol B | 4,4'-(butane-2,2-diyl)diphenol |
| 1571-75-1 | bisphenol AP | 4,4'-(1-phenylethane-1,1-diyl)diphenol |
| 5613-46-7 | tetramethyl bisphenol A | 4,4'-(propane-2,2-diyl)bis(2,6-dimethylphenol) |
| 79-97-0 | 2,2-bis(4-hydroxy-3-methylphenyl)propane | 4,4'-(propane-2,2-diyl)bis(2-methylphenol) |
| 27955-94-8 | 1,1',1''-tris(4-hydroxyphenyl)ethane | 4,4',4''-(ethane-1,1,1-triyl)triphenol |
| 599-64-4 | 4-α-cumyl phenol | 4-(2-phenylpropan-2-yl)phenol |
| 2167-51-3 | bisphenol P | 4,4'-(1,4-phenylenebis(propane-2,2-diyl))diphenol |
| 14868-03-2 | bisphenol C | 4,4'-(2,2-dichloroethene-1,1-diyl)diphenol |
| 80-05-7 | bisphenol A | 4,4'-(propane-2,2-diyl)diphenol |
| 70-30-4 | hexachlorophene | 6,6'-methylenebis(2,4,5-trichlorophenol) |
| 110726-28-8 | $\alpha,\alpha,\alpha'$-tris(4-hydroxyphenyl)-1-ethyl-4-isopropylbenzene | 4,4'-(1-(4-(2-(4-hydroxyphenyl)propan-2-yl)phenyl)ethane-1,1-diyl)diphenol |
| 2716-10-1 | $\alpha,\alpha'$-bis(4-aminophenyl)-1,4-diisopropylbenzene | 4,4'-(1,4-phenylenebis(propane-2,2-diyl))dianiline |
| 57100-74-0 | 2,2-bis(3-cyclohexyl-4-hydroxyphenyl)propane | 4,4'-(propane-2,2-diyl)bis(2-cyclohexylphenol) |
| 24038-68-4 | 2,2-bis(2-hydroxy-5-biphenyl)propane | 5,5''-(propane-2,2-diyl)bis([1,1'-biphenyl]-2-ol)) |
| 1675-54-3 | 2,2-bis(4-glycidyloxyphenyl)propane | 2,2'-(((propane-2,2-diylbis(4,1-phenylene))bis(oxy))bis(methylene))bis(oxirane) |
| 10192-62-8 | bisphenol A diacetate | propane-2,2-diylbis(4,1-phenylene) diacetate |
| 4162-45-2 | tetrabromobisphenol A bis(2-hydroxyethyl)ether | 2,2'-((propane-2,2-diylbis(2,6-dibromo-4,1-phenylene))bis(oxy))diethanol |
| 127-54-8 | 2,2-bis(4-hydroxy-3-isopropylphenyl)propane | 4,4'-(propane-2,2-diyl)bis(2-isopropylphenol) |
| 3539-42-2 | 4,4'-isopropylidenediphenoxyacetic acid | 2,2'-((propane-2,2-diylbis(4,1-phenylene))bis(oxy))diacetic acid |
| 13080-86-9 | 2,2-bis[4-(4-aminophenoxy)-phenyl]propane | 4,4'-((propane-2,2-diylbis(4,1-phenylene))bis(oxy))dianiline |
| 36395-57-0 | $\alpha,\alpha'$-bis(4-hydroxy-3,5-dimethylphenyl)-1,4-diisopropylbenzene | 4,4'-(1,4-phenylenebis(propane-2,2-diyl))bis(2,6-dimethylphenol) |
| 2024-88-6 | 2,2-bis(4-chloroformyloxyphenyl)propane | propane-2,2-diylbis(4,1-phenylene) dicarbonochloridate |
| 32113-46-5 | 2,2-bis(3-sec-butyl-4-hydroxyphenyl)propane | 2-butan-2-yl-4-[2-(3-butan-2-yl-4-hydroxyphenyl)propan-2-yl]phenol |
| 620-92-8 | bisphenol F | 4,4'-methylenediphenol |
| 84-16-2 | hexestrol | 4,4'-(hexane-3,4-diyl)diphenol |
| 1156-51-0 | 2,2-bis(4-cyanatophenyl)propane | 4,4'-(propane-2,2-diyl)bis(cyanatobenzene) |
| 479-13-0 | coumestrol | 3,9-dihydroxy-6H-benzofuro[3,2-c]chromen-6-one |
| 1415-73-2 | barbaloin | (10*S*)-1,8-dihydroxy-3-(hydroxymethyl)-10-[(2*S*,3*R*,4*R*,5*S*,6*R*)-3,4,5-trihydroxy-6-(hydroxymethyl)oxan-2-yl]-10*H*-anthracen-9-one |
| 961-29-5 | isoliquirtigenin | (*E*)-1-(2,4-dihydroxyphenyl)-3-(4-hydroxyphenyl)prop-2-en-1-one |
| 2467-25-6 | 4,4'-methylenebis(2-methylphenol) | 4,4'-methylenebis(2-methylphenol) |
| 17345-66-3 | 2,3,4-trihydroxydiphenylmethane | 4-benzylbenzene-1,2,3-triol |
| 75804-28-3 | 2,3-dimethyl-2,3-butanediamine | 2,3-dimethylbutane-2,3-diamine |
| 2081-08-5 | bisphenol E | 4,4'-(ethane-1,1-diyl)diphenol |
| 2971-36-0 | HPTE | 4,4'-(2,2,2-trichloroethane-1,1-diyl)diphenol |
| 83558-87-6 | 2,2-bis(3-amino-4-hydroxyphenyl)hexafluoropropane | 4,4'-(perfluoropropane-2,2-diyl)bis(2-aminophenol) |
| 47250-53-3 | 2,2-bis(3-aminophenyl)hexafluoropropane | 3,3'-(perfluoropropane-2,2-diyl)dianiline |
| 116325-74-7 | 2,2-bis(3-amino-4-methylphenyl)hexafluoropropane | 5,5'-(perfluoropropane-2,2-diyl)bis(2-methylaniline) |
| 1095-78-9 | 2,2-bis(4-aminophenyl)hexafluoropropane | 4,4'-(perfluoropropane-2,2-diyl)dianiline |
| 69563-88-8 | 2,2-bis[4-(4-aminophenoxy)phenyl]hexafluoropropane | 4,4'-(((perfluoropropane-2,2-diyl)bis(4,1-phenylene))bis(oxy))dianiline |
| 1478-61-1 | bisphenol AF | 4,4'-(perfluoropropane-2,2-diyl)diphenol |
| 1107-00-2 | 4,4'-(hexafluoroisopropylidene)diphthalic anhydride | 5,5'-(perfluoropropane-2,2-diyl)bis(isobenzofuran-1,3-dione) |
| 1171-47-7 | 2,2-bis(4-carboxyphenyl)hexafluoropropane | 4,4'-(perfluoropropane-2,2-diyl)dibenzoic acid |
| 10224-18-7 | 2,2-bis(4-isocyanatophenyl)hexafluoropropane | 4,4'-(perfluoropropane-2,2-diyl)bis(isocyanatobenzene) |
| 83558-76-3 | hexafluoro-2,2-diphenylpropane | (perfluoropropane-2,2-diyl)dibenzene |
| 4221-68-5 | 1,1-bis(3-cyclohexyl-4-hydroxyphenyl)cyclohexane | 4,4'-(cyclohexane-1,1-diyl)bis(2-cyclohexylphenol) |
| 15499-84-0 | 9,9-bis(4-aminophenyl)fluorene | 4,4'-(9H-fluorene-9,9-diyl)dianiline |
| 184355-68-8 | 4,4'-(2-hydroxybenzylidene)-bis(2,3,6-trimethylphenol) | 4,4'-((2-hydroxyphenyl)methylene)bis(2,3,6-trimethylphenol) |
| 6807-17-6 | 4,4'-(1,3-dimethylbutylidene)diphenol | 4,4'-(4-methylpentane-2,2-diyl)diphenol |
| 3236-71-3 | 9,9-bis(4-hydroxyphenyl)fluorene | 4,4'-(9H-fluorene-9,9-diyl)diphenol |
| 88938-12-9 | 9,9-bis(4-hydroxy-3-methylphenyl)fluorene | 4,4'-(9H-fluorene-9,9-diyl)bis(2-methylphenol) |
| 74462-02-5 | 4,4'-(2-ethylhexylidene)diphenol | 4,4'-(2-ethylhexane-1,1-diyl)diphenol |
| 117344-32-8 | 9,9-bis[4-(2-hydroxyethoxy)phenyl]fluorene | 2,2'-(((9H-fluorene-9,9-diyl)bis(4,1-phenylene))bis(oxy))diethanol |
| 2362-14-3 | 1,1-bis(4-hydroxy-3-methylphenyl)cyclohexane | 4,4'-(cyclohexane-1,1-diyl)bis(2-methylphenol) |
| 3282-99-3 | 1,1-bis(4-aminophenyl)cyclohexane | 4,4'-(cyclohexane-1,1-diyl)dianiline |
| 843-55-0 | bisphenol Z | 4,4'-(cyclohexane-1,1-diyl)diphenol |
| 13595-25-0 | 1,3-bis[2-(4-hydroxyphenyl)-2-propyl]benzene | 4,4'-(1,3-phenylenebis(propane-2,2-diyl))diphenol |
| 20601-38-1 | 4,4'-bicyclohexanol | [1,1'-bi(cyclohexane)]-4,4'-diol |
| 1980-4-69 | 2,2-bis(4-hydroxycyclohexyl)propane | 4,4'-(propane-2,2-diyl)dicyclohexanol |
| 2433-14-6 | 4-cyclohexylcyclohexanol | [1,1'-bi(cyclohexan)]-4-ol |
| 119-42-6 | 2-cyclohexylphenol | 2-cyclohexylphenol |
| 947-42-2 | diphenylsilanediol | diphenylsilanediol |
| 20714-70-9 | 4-(phenylazo)phenol | (E)-4-(phenyldiazenyl)phenol |
| 501-36-0 | resveratrol | (E)-5-(4-hydroxystyryl)benzene-1,3-diol |
| 3127-14-8 | spirobiromane | 4,4,4',4'-tetramethyl-2,2'-spirobi[chroman]-7,7'-diol |
| 2246-46-0 | 4-(2-Thiazolylazo)resorcinol | (E)-4-(thiazol-2-yl diazenyl)benzene-1,3-diol |
| 32737-35-2 | 6,6',7,7'-tetrahydroxy-4,4,4',4'-tetramethyl-2,2'-spirobichroman | 4,4,4',4'-tetramethyl-2,2'-spirobi[chroman]-6,6',7,7'-tetraol |
| 269409-97-4 | 4-(4,4,5,5-Tetramethyl-1,3,2-dioxaborolan-2-yl)phenol | 2-(4,4,5,5-tetramethyl-1,3,2-dioxaborolan-2-yl)phenol |
| 611-99-4 | 4,4'-dihydroxybenzophenone | bis(4-hydroxyphenyl)methanone |
| 90-96-0 | 4,4'-dimethoxybenzophenone | bis(4-methoxyphenyl)methanone |
| 61445-50-9 | 2,3',4,4'-tetrahydroxybenzophenone | (2,4-dihydroxyphenyl)(3,4-dihydroxyphenyl)methanone |
| 131-55-5 | 2,2',4,4'-tetrahydroxybenzophenone | bis(2,4-dihydroxyphenyl)methanone |
| 345-92-6 | 4,4'-difluorobenzophenone | bis(4-fluorophenyl)methanone |
| 131-54-4 | 2,2'-dihydroxy-4,4'-dimethoxybenzophenone | bis(2-hydroxy-4-methoxyphenyl)methanone |
| 90-98-2 | 4,4'-dichlorobenzophenone | bis(4-chlorophenyl)methanone |
| 2421-28-5 | 3,3',4,4'-benzophenonetetracarboxylic dianhydride | 5,5'-carbonylbis(isobenzofuran-1,3-dione) |
| 85-58-5 | benzophenone-2,4'-dicarboxylic acid monohydrate | 2-(4-carboxybenzoyl)benzoic acid |
| 342-25-6 | 2,4'-difluorobenzophenone | (2-fluorophenyl)(4-fluorophenyl)methanone |
| 611-98-3 | 4,4'-diaminobenzophenone | bis(4-aminophenyl)methanone |
| 83846-85-9 | 4-benzoyl 4'-methyldiphenyl sulfide | phenyl(4-(p-tolylthio)phenyl)methanone |
| 964-68-1 | benzophenone-4,4'-dicarboxylic acid | 4,4'-carbonyldibenzoic acid |
| 131-53-3 | 2,2'-dihydroxy-4-methoxybenzophenone | (2-hydroxy-4-methoxyphenyl)(2-hydroxyphenyl)methanone |
| 85-29-0 | 2,4'-dichlorobenzophenone | (2-chlorophenyl)(4-chlorophenyl)methanone |
| 1470-79-7 | 2,4,4'-trihydroxybenzophenone | (2,4-dihydroxyphenyl)(4-hydroxyphenyl)methanone |
| 835-11-0 | 2,2'-dihydroxybenzophenone | bis(2-hydroxyphenyl)methanone |
| 21222-05-9 | 3,3'-dinitrobenzophenone | bis(3-nitrophenyl)methanone |
| 611-79-0 | 3,3'-diaminobenzophenone | bis(3-aminophenyl)methanone |
| 2958-36-3 | 2-amino-2',5-dichlorobenzophenone | (2-amino-5-chlorophenyl)(2-chlorophenyl)methanone |
| 33077-87-1 | 2,2',4-trimethoxybenzophenone | (2,4-dimethoxyphenyl)(2-methoxyphenyl)methanone |
| 3708-39-2 | 4,4'-bis(methylamino)benzophenone | bis(4-(methylamino)phenyl)methanone |
| 119-61-9 | benzophenone | benzophenone |
| 31127-54-5 | 2,3,4,4'-tetrahydroxybenzophenone | (4-hydroxyphenyl)(2,3,4-trihydroxyphenyl)methanone |
| 118-82-1 | 4,4'-methylenebis(2,6-di-tert-butylphenol) | 4,4'-methylenebis(2,6-di-tert-butylphenol) |
| 122-25-8 | methylenedisalicylic acid | 5,5'-methylenebis(2-hydroxybenzoic acid) |
| 105391-33-1 | bis(3-ethyl-5-methyl-4-maleimidophenyl)methane | 1,1'-(methylenebis(2-ethyl-6-methyl-4,1-phenylene))bis(1H-pyrrole-2,5-dione) |
| 97-23-4 | 2,2'-methylenebis(4-chlorophenol) | 2,2'-methylenebis(4-chlorophenol) |
| 13676-54-5 | 4,4'-bismaleimidodiphenylmethane | 1,1'-(methylenebis(4,1-phenylene))bis(1H-pyrrole-2,5-dione) |
| 19900-72-2 | 4,4'-methylenebis(2-ethyl-6-methylaniline) | 4,4'-methylenebis(2-ethyl-6-methylaniline) |
| 838-88-0 | 4,4'-diamino-3,3'-dimethyldiphenylmethane | 4,4'-methylenebis(2-methylaniline) |
| 101-77-9 | 4,4'-diaminodiphenylmethane | 4,4'-methylenedianiline |
| 101-61-1 | Bis[4-dimethylamino-phenyl]methane | 4,4'-methylenebis(N,N-dimethylaniline) |
| 88-24-4 | 2,2'-methylenebis(6-tert-butyl-4-ethylphenol) | 6,6'-methylenebis(2-(tert-butyl)-4-ethylphenol) |
| 101-14-4 | 4,4'-methylenebis(2-chloroaniline) | 4,4'-methylenebis(2-chloroaniline) |
| 5384-21-4 | 4,4'-methylenebis(2,6-dimethylphenol) | 4,4'-methylenebis(2,6-dimethylphenol) |
| 19430-83-2 | 3,4'-diaminodiphenylmethane | 3-(4-aminobenzyl)aniline |
| 119-47-1 | 2,2'-methylenebis(6-tert-butyl-p-cresol) | 6,6'-methylenebis(2-(tert-butyl)-4-methylphenol) |
| 1817-74-9 | 4,4'-dinitrodiphenylmethane | bis(4-nitrophenyl)methane |
| 42240-73-3 | Bis(4-amino-2,3-dichlorophenyl)methane | 4,4'-methylenebis(2,3-dichloroaniline) |
| 3236-63-3 | 2,2'-methylenebis(4-methylphenol) | 2,2'-methylenebis(4-methylphenol) |
| 2467-03-0 | 2,4'-dihydroxydiphenylmethane | 2-(4-hydroxybenzyl)phenol |
| 457-68-1 | 4,4'-difluorodiphenylmethane | bis(4-fluorophenyl)methane |
| 101-68-8 | 4,4'-diphenylmethane diisocyanate, (4,4'-Methylenebis(phenyl Isocyanate)) | bis(4-isocyanatophenyl)methane |
| 139-25-3 | 4,4'-diisocyanato-3,3'-dimethyldiphenylmethane | bis(4-isocyanato-3-methylphenyl)methane |
| 19471-12-6 | 3,3'-diaminodiphenylmethane | 3,3'-methylenedianiline |
| 2467-02-9 | 2,2'-dihydroxydiphenylmethane | 2,2'-methylenediphenol |
| 1844-01-5 | 4,4'-dihydroxytetraphenylmethane | 4,4'-(diphenylmethylene)diphenol |
| 174462-43-2 | 2,3,4,4'-tetrahydroxydiphenylmethane | 4-(4-hydroxybenzyl)benzene-1,2,3-triol |

**Table S2.**
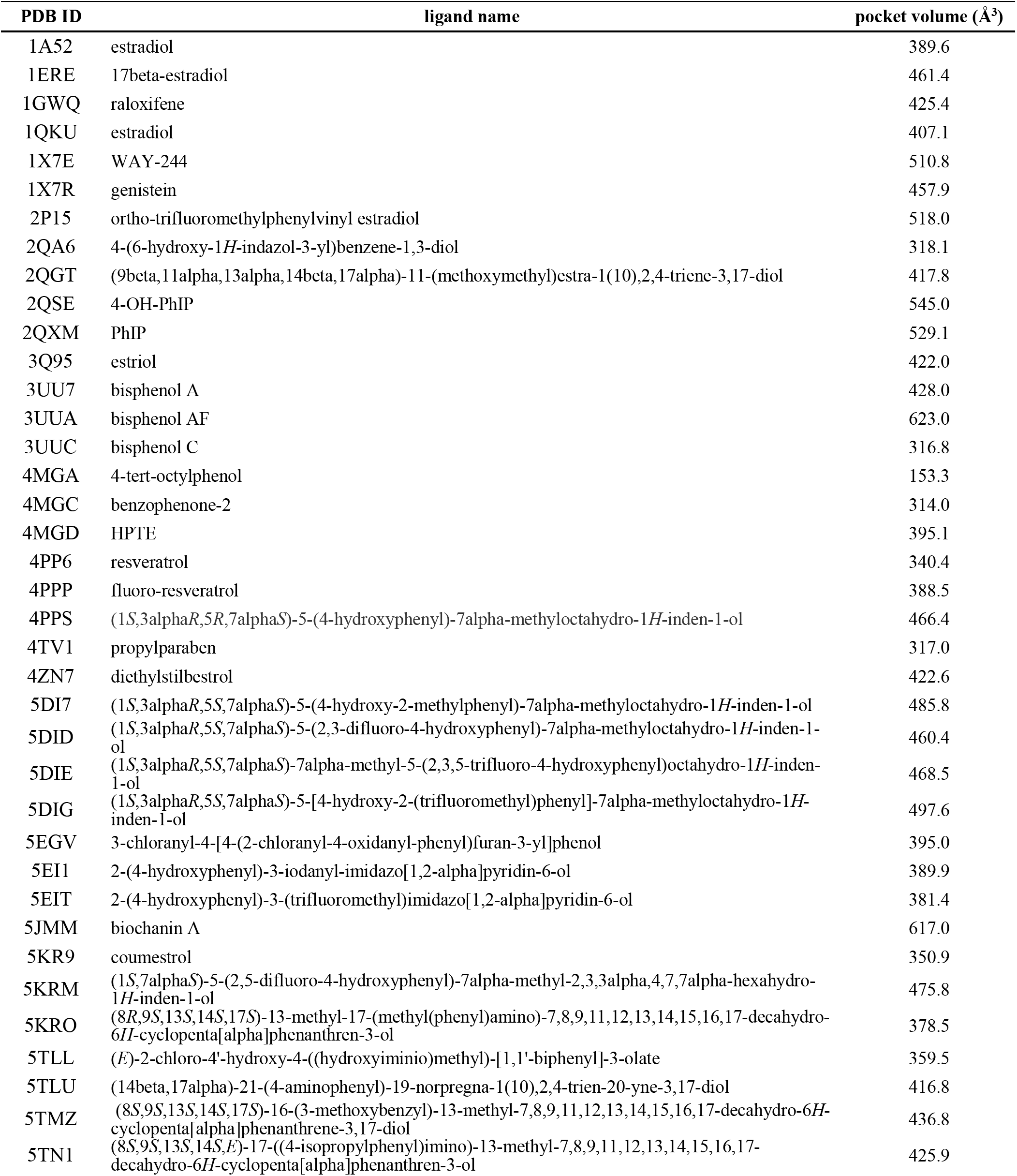
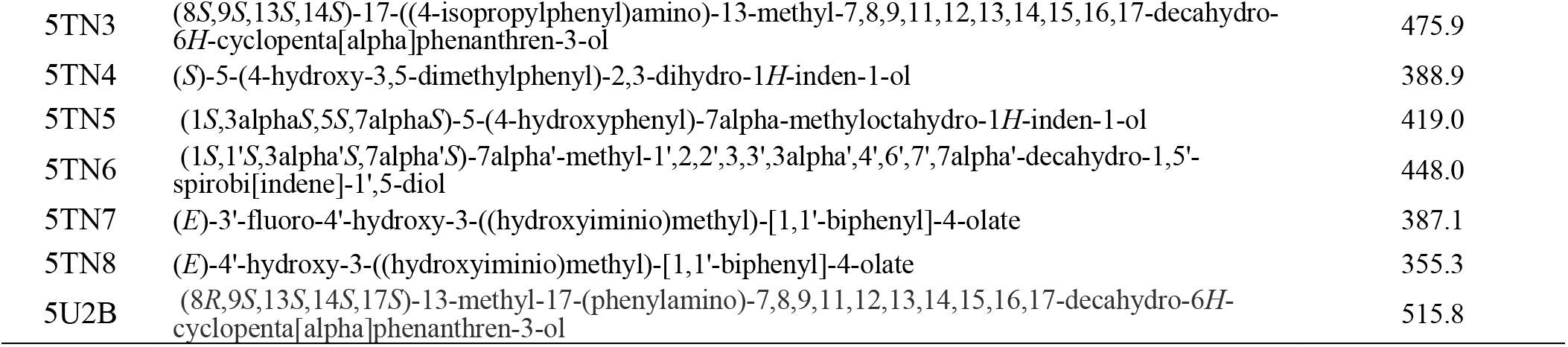
PDB IDs of ERα utilized for calculating of the volumes of each ligand-binding pocket. PDB IDs are listed in alphabetical order.

**Table S3.** PDB IDs of ERβ agonist structures utilized for calculating of the volumes of each ligand binding pocket. PDB IDs are listed in alphabetical order.

| PDB ID | ligand name | pocket volume (Å<sup>3</sup>) |
| --- | --- | --- |
| 1QKN | raloxifene | 382.3 |
| 1U3Q | 4-(6-hydroxy-benzo[delta]isoxazol-3-yl)benzene-1,3-diol | 344.9 |
| 1U3R | 2-(5-hydroxy-naphthalen-1-yl)-1,3-benzoxazol-6-ol | 364.3 |
| 1U3S | 3-(6-hydroxy-naphthalen-2-yl)-benzo[delta]isoxazol-6-ol | 407.9 |
| 1X76 | 5-hydroxy-2-(4-hydroxyphenyl)-1-benzofuran-7-carbonitrile | 354.4 |
| 1X78 | [5-hydroxy-2-(4-hydroxyphenyl)-1-benzofuran-7-yl]acetonitrile | 349.4 |
| 1X7B | 2-(3-fluoro-4-hydroxyphenyl)-7-vinyl-1,3-benzoxazol-5-ol | 314.0 |
| 1X7J | genistein | 375.9 |
| 1YY4 | 1-chloro-6-(4-hydroxyphenyl)-2-naphthol | 408.5 |
| 1YYE | 3-(3-fluoro-4-hydroxyphenyl)-7-hydroxy-1-naphthonitrile | 402.6 |
| 1ZAF | 3-bromo-6-hydroxy-2-(4-hydroxyphenyl)-1*H*-inden-1-one | 503.8 |
| 2J7X | estradiol | 423.0 |
| 2J7Y | (16alpha,17alpha)-estra-1,3,5(10)-triene-3,16,17-triol | 372.6 |
| 2NV7 | 4-(4-hydroxyphenyl)-1-naphthaldehyde oxime | 340.6 |
| 2YJD | 4-(2-propan-2-yloxybenzimidazol-1-yl)phenol | 468.1 |
| 2YLY | *n*-cyclopropyl-4-oxidanyl-*n*-[(2*R*)-2-oxidanyl-2-phenyl-propyl]benzenesulfonamide | 226.3 |
| 3OLL | estradiol | 361.9 |
| 3OLS | estradiol | 404.9 |
| 3OMO | 2-(trifluoroacetyl)-1,2,3,4-tetrahydroisoquinolin-6-ol | 289.0 |
| 3OMP | 2-(trifluoroacetyl)-1,2,3,4-tetrahydroisoquinolin-7-ol | 321.3 |
| 3OMQ | 2-[(trifluoromethyl)sulfonyl]-1,2,3,4-tetrahydroisoquinolin-6-ol | 346.9 |
| 4J24 | estradiol | 324.9 |
| 4J26 | estradiol | 434.6 |
| 4ZI1 | 2-(4-hydroxyphenyl)-7-methyl-3-phenyl-1*H*-inden-5-ol | 350.4 |
| 5TOA | estradiol | 361.0 |

**Table S4.** PDB IDs of ERβ LBDs utilized for SiteFinder calculations to analyze ligand-binding sites. PDB IDs are listed in alphabetical order.

| PDB ID | rank of 1st site | rank of 2nd site | ligand | active or inactive | position of H12* |
| --- | --- | --- | --- | --- | --- |
| 1HJ1 | 1 | 7 | ICI164,384 | inactive | free |
| 1L2J | 1 | none | (*R,R*)-5,11-cis-diethyl-5,6,11,12-tetrahydrochrysene-2,8-diol | inactive | CBS |
| 1NDE | 1 | none | 4-(2-([4-([3-(4-chlorophenyl)propyl)sulfanyl]-6-(1-piperazinyl)-1,3,5-triazin-2-yl]amino)ethyl)phenol | inactive | CBS |
| 1QKM | 1 | 9 | genistein | inactive | free |
| 1QKN | 1 | 10 | raloxifene | inactive | free |
| 1U3Q | 1 | none | 4-(6-hydroxy-benzo[d]isoxazol-3-yl)benzene-1,3-diol | active | active position |
| 1U3R | 1 | none | 2-(5-hydroxy-naphthalen-1-yl)-1,3-benzoxazol-6-ol | active | active position |
| 1U3S | 1 | none | 3-(6-hydroxy-naphthalen-2-yl)-benzo[d]isoxazol-6-ol | active | active position |
| 1U9E | 1 | 3 | 2-(4-hydroxy-phenyl)benzofuran-5-ol | active | active position |
| 1X76 | 1 | 3 | 5-hydroxy-2-(4-hydroxyphenyl)-1-benzofuran-7-carbonitrile | active | active position |
| 1X78 | 1 | none | [5-hydroxy-2-(4-hydroxyphenyl)-1-benzofuran-7-yl]acetonitrile | active | active position |
| 1X7B | 1 | none | 2-(3-fluoro-4-hydroxyphenyl)-7-vinyl-1,3-benzoxazol-5-ol | active | active position |
| 1X7J | 5 | none | genistein | active | active position |
| 1YY4 | 1 | none | 1-chloro-6-(4-hydroxyphenyl)-2-naphthol | active | active position |
| 1YYE | 1 | 17 | 3-(3-fluoro-4-hydroxyphenyl)-7-hydroxy-1-naphthonitrile | active | active position |
| 1ZAF | 1 | 3 | 3-bromo-6-hydroxy-2-(4-hydroxyphenyl)-1*H*-inden-1-one | active | active position |
| 2FSZ | 5 | 11 | 4-hydroxytamoxifen | inactive | free |
| 2GIU | 1 | none | (9a*S*)-4-bromo-9a-butyl-7-hydroxy-1,2,9,9a-tetrahydro-3*H*-fluoren-3-one | inactive | CBS |
| 2I0G | 1 | none | (3a*S*,4*R*,9b*R*)-4-(4-hydroxyphenyl)-1,2,3,3a,4,9b-hexahydrocyclopenta[c]chromen-8-ol | inactive | CBS |
| 2I0J | 1 | 9 | (3a*S*,4*R*,9b*R*)-4-(4-hydroxyphenyl)-1,2,3,3a,4,9b-hexahydrocyclopenta[c]chromen-8-ol | inactive | free |
| 2J7X | 1 | none | estradiol | active | active position |
| 2J7Y | 1 | none | (16alpha,17alpha)-estra-1,3,5(10)-triene-3,16,17-triol | active | active position |
| 2JJ3 | 1 | none | (3a*S*,4*R*,9b*R*)-4-(4-hydroxyphenyl)-6-(methoxymethyl)-1,2,3,3a,4,9b-hexahydrocyclopenta[c]chromen-8-ol | inactive | CBS |
| 2NV7 | 1 | 13 | 4-(4-hydroxyphenyl)-1-naphthaldehyde oxime | active | active position |
| 2POG | 1 | 8 | (3a*S*,4*R*,9b*R*)-4-(4-hydroxyphenyl)-1,2,3,3a,4,9b-hexahydrocyclopenta[c]chromen-9-ol | inactive | free |
| 2QTU | 1 | none | (3a*S*,4*R*,9b*R*)-2,2-difluoro-4-(4-hydroxyphenyl)-6-(methoxymethyl)-1,2,3,3a,4,9b-hexahydrocyclopenta[c]chromen-8-ol | inactive | CBS |
| 2YJD | 1 | 3 | 4-(2-propan-2-yloxybenzimidazol-1-yl)phenol | active | active position |
| 2YLY | 2 | 5 | *N*-cyclopropyl-4-oxidanyl-*N*-[(2*R*)-2-oxidanyl-2-phenylpropyl]benzenesulfonamide | active | active position |
| 2Z4B | 1 | none | (3a*S*,4*R*,9b*R*)-2,2-difluoro-4-(4-hydroxyphenyl)-1,2,3,3a,4,9b-hexahydrocyclopenta[c]chromen-8-ol | inactive | CBS |
| 3OLL | 1 | 8 | estradiol | active | active position |
| 3OLS | 1 | none | estradiol | active | active position |
| 3OMO | 1 | none | 2-(trifluoroacetyl)-1,2,3,4-tetrahydroisoquinolin-6-ol | active | active position |
| 3OMP | 1 | 10 | 2-(trifluoroacetyl)-1,2,3,4-tetrahydroisoquinolin-7-ol | active | active position |
| 3OMQ | 1 | none | 2-[(trifluoromethyl)sulfonyl]-1,2,3,4-tetrahydroisoquinolin-6-ol | active | active position |
| 4J24 | 1 | 10 | estradiol | active | active position |
| 4J26 | 1 | 9 | estradiol | active | active position |
| 4ZI1 | 1 | none | 2-(4-hydroxyphenyl)-7-methyl-3-phenyl-1*H*-inden-5-ol | active | active position |
| 5TOA | 1 | 7 | estradiol | active | active position |

\* “CBS” means that H12 is located in an inactivated position on the ERβ coactivator-binding site (CBS); 'free' means helix 12 is not visualized or is far outside of the LBD.

**Table S5.** Compounds names, CAS RN, and ligand IDs of CDS-Core or Chemical IDs from the Protein Data Bank (PDB); 3D coordinates were utilized for docking simulation experiments. Chemical IDs from PDB are designated by three letters.

|  | common names | CAS RN® | Ligand ID |
| --- | --- | --- | --- |
| 1 | bisphenol C | 14868-03-2 | 0D1 (PDB) |
| 2 | 4,4'-(1,3-dimethylbutylidene)bisphenol | 6807-17-6 | ZUHRAX |
| 3 | 2,2-bis(p-hydroxyphenyl)-1,1,1- trichloroethane (HPTE) | 2971-36-0 | - |
| 4 | bisphenol AF | 1478-61-1 | TIBVOQ |
| 5 | bisphenol Z | 843-55-0 | - |
| 6 | 4,4'-(2-ethylhexylidene)bisphenol | 74462-02-5 | - |
| 7 | 4,4'-(2-hydroxybenzylidene)-bis(2,3,6-trimethylphenol) | 184355-68-8 | - |
| 8 | bisphenol B | 77-40-7 | - |
| 9 | 1,1-bis(4-hydroxy-3-methylphenyl)cyclohexane | 2362-14-3 | SIJHOJ |
| 10 | bisphenol M | 13595-25-0 | - |
| 11 | bisphenol AP | 1571-75-1 | - |
| 12 | $\alpha, \alpha, \alpha'$-tris(4-hydroxyphenyl)-1-ethyl-4-isopropylbenzene | 110726-28-8 | - |
| 13 | 2,2-bis(3-amino-4-hydroxyphenyl)hexafluoropropane | 83558-87-6 | - |
| 14 | 9,9-Bis(4-hydroxyphenyl)fluorene | 3236-71-3 | ABUCOP |
| 15 | 9,9-bis(4-hydroxy-3-methylphenyl)fluorene | 15499-84-0 | XOGJEI |
| 16 | bisphenol P | 2167-51-3 | - |
| 17 | 2,2-bis[4-(4-aminophenoxy)phenyl]hexafluoropropane | 69563-88-8 | HOYZOL |
| 18 | 2,2-bis(4-hydroxy-3-methylphenyl)propane | 79-97-0 | REGKOF |
| 19 | bisphenol A | 80-05-7 | 2OH (PDB) |
| 20 | $\alpha, \alpha'$-bis(4-hydroxy-3,5-dimethylphenyl)-1,4-diisopropylbenzene | 36395-57-0 | ACAYIN |

